# A multi-model approach to assessing local and global cryo-EM map quality

**DOI:** 10.1101/128561

**Authors:** Mark A. Herzik, James S. Fraser, Gabriel C. Lander

## Abstract

There does not currently exist a standardized indicator of how well a cryo-EM-derived model represents the density from which it was generated. We present a straightforward methodology that utilizes freely available tools to generate a suite of independent models and to evaluate their convergence in an EM density. These analyses provide both a quantitative and qualitative assessment of the precision of the models and their representation of the density, respectively, while concurrently providing a platform for assessing both global and local EM map quality. We further use standardized datasets to provide an expected model–model agreement criterion for EM maps reported to be at 5 Å resolution or better. Associating multiple atomic models with a deposited EM map provides a rapid and accessible reporter of convergence, a strong indicator of highly resolved molecular detail, and is an important step toward an FSC-independent assessment of map and model quality.

## Introduction

In order to perform important cellular processes, biological macromolecules adopt a variety of conformations, each with the potential to serve a distinct functional role (McCammon 1977, Frauenfelder 1991, Alberts 1998). Molecular motions between these conformations are integral to biological function, and a molecular-level description of a macromolecule’s conformational landscape is critical to understanding its role in a cellular context (Smock and Gierasch 2009). Cryo-electron microscopy (cryo-EM) proves to be an increasingly powerful structural technique for studying conformationally or compositionally heterogeneous macromolecules. Both types of heterogeneity are preserved upon vitrification during cryo-EM sample generation and can be discerned structurally using “*in silico* purification” that leverages sophisticated 2D and 3D classification and refinement protocols to identify homogenous species (Sigworth 2007, Nogales and Scheres 2015). Indeed, a single data collection often yields several conformationally (Bai 2015, Zhou 2015, Abeyrathne 2016, Banerjee 2016, He 2016) or compositionally distinct structures (Shen 2015, Davis 2016, Tsai 2017). Although image-processing strategies typically strive to identify the most homogenous particles within a dataset, cryo-EM reconstructions frequently possess large internal variations in resolution. This variability in resolution presumably results from residual structural heterogeneity (encompassing both compositional and conformational) and imperfect particle alignment during the reconstruction.

Although cryo-EM reconstructions exhibit local resolution variations, a single, static atomic model is typically used to represent these structures, an approach inherited from the X-ray crystallography community. As such, a single-conformer model at unit occupancy is presented, with uncertainty modeled by an isotropic Gaussian distribution of the position of each atom (atomic displacement parameter, or B-factor). The B-factor is an incomplete description of the extent of heterogeneity exhibited by a structure because isotropic motion is a poor approximation for most protein conformational heterogeneity and it additionally convolves uncertainty, model error, and refinement restraints (Kuriyan 1986). The local accuracy and precision a single static model cannot be evaluated by any single metric, forcing users to examine a combination of B-factors, improbable geometric outliers, and local correlation to the density (X-ray or EM) as a means to assess model quality. However, the relationship between the atomic model and the experimental data differ greatly between X-ray crystallography and EM. Whereas atomic models derived from X-ray diffraction are inextricably linked to the data due to the fact that iterative improvements in phase information arise from an improved atomic model, EM potential density maps are generated independently of an atomic model. Because the target density remains unchanged during atomic model generation and refinement in EM, the question of how well a model converges is less subject to model bias and primarily reflects the local resolution/quality of the map.

Many alternative representations of macromolecular structure have been proposed previously that attempt to overcome the limitations of using a single conformer and B-factors (reviewed in (Woldeyes 2014)). In X-ray crystallography, these alternative representations can be grouped into two different classes: 1) an ensemble of multiple complete models that each are presented as *independent* but equally valid interpretations of the structural data (DePristo 2004) and 2) ensembles of multiple conformations that are presented as a *collective* interpretation of the structural data (Levin 2007, Burnley 2012, Keedy 2015). These methods are similar to the current practices for generating NMR ensembles that are either equally valid interpretations of restraints (Wuthrich 1990) or that use more sophisticated averaging schemes (Lange 2008), both of which are routinely deposited at the Protein Data Bank (www.rcsb.org) (Berman 2000). The conformational heterogeneity represented by these ensembles arises not only from the dynamic nature of the macromolecule, but also from uncertainties in image processing and model generation/refinement. While the collective ensembles can, in principle, accurately capture discretely conformations observable at high resolution, the independent ensembles are biased towards the most populated conformation and therefore yield an estimate of precision (Terwilliger 2007). However, both model types demonstrate the limitations of the single static model and the inadequacy of the B-factor approximation to model the underlying conformational hetereogeneity (Kuzmanic 2014).

Despite proposals that the fields adopt alternative multi-model or multi-conformer representations in X-ray crystallography (Furnham 2006), the practice has not been widely adopted. However, because independent models provide a reasonable local estimate of the lower-bound of model precision and do not introduce additional parameters (Terwilliger 2007), they may offer additional utility in addressing an emerging challenge in EM: evaluating local map quality and the uncertainty of the model. This issue emerges because EM lacks foolproof and robust measures of local and global resolution. Whereas X-ray crystallography has a stringent set of metrics that can be used to assess the global quality of diffraction data, the cryo-EM field primarily relies on the “gold standard” Fourier Shell Correlation (FSC) (Henderson 2012, Scheres and Chen 2012). The FSC reports on the resolution of a determined structure by measuring the correlation between two independently refined “half-maps” across spatial frequencies (Saxton and Baumeister 1982, van Heel and Harauz 1986, Grigorieff 2000, Penczek 2010, Henderson 2012, Scheres and Chen 2012), a measure of the self-consistency of the data rather than a true measure of “resolution”. Reporting the resolution of a structure according to the FSC is limited, as there are a variety of ways in which the global resolution can be over-estimated, such as by excluding poorly resolved regions from calculation of the FSC through application of a 3D mask. Furthermore, reconstructions with preferred orientation can contain extensive anisotropy in resolution that will not be evident in the FSC curve (Penczek 2002, Lander 2013, Urnavicius 2015). This problem is further compounded by the fact that a single value from this curve is generally ascribed to an EM structure, providing only an approximation of the quality of the molecular details contained within a reconstruction, and without indication of the degree of resolution variation. As a result, care must be taken when evaluating a map solely on the FSC-reported resolution value, as this does not guarantee map quality or model correctness.

Currently, someone downloading EM-derived atomic coordinates from the PDB can rely mainly on the B-factor for indication of how well a model fits the EM density. Because the B-factor convolves model error, refinement restraints, and conformational variability, it can be a misleading metric of modeling precision (Kuzmanic 2014). In contrast, the ability of multiple independently generated atomic coordinates to converge on a single point is a strong indicator of highly resolved molecular detail, and we wish to use this concept to assign a “convergence factor” to every residue in a structure, as well as present the user with an intuitive and physically meaningful representation of this factor. Associating multiple atomic models with a deposited EM map provides a rapid and accessible reporter of convergence, and is an important supplement to the traditional FSC-based assessment of map and model quality.

Here we present a straightforward methodology that utilizes freely available tools to provide metrics for both global and local EM map quality, while concurrently providing a platform to assess the reported EM map quality independently of FSC. We show that the generation of multiple independent models using an EM density and subsequent analysis of their atomic agreement statistics provide model quality metrics that directly correlate with global and local map quality. In addition, this multi-model approach provides both a quantitative and qualitative assessment of the precision of the models and their representation of the density, respectively. We further show that this analysis can be applied to most EM maps reported to be at 5 Å resolution or better, and use standardized *in silico* datasets to provide an expected multi-model agreement criterion for EM maps across a broad resolution range.

## Results

The main goal of this study is to test the ability of multiple models that occupy unique coordinate space to converge to a single uniform set of atomic coordinates, given the molecular envelope of a cryo-EM reconstruction as a restraint. For a given set of structurally distinct atomic models that have been refined into an EM density, the RMSD of each C_α_ can be used as a metric of model convergence, serving as a lower bound of the coordinate precision of an atomic ensemble (Terwilliger 2007). Presumably, the convergence of a given set of distinct coordinates will correspond to the quality of the density. Higher resolution structural elements, where side-chain densities are clearly resolved, should produce RMSDs close to zero, and displacements should be accurately modeled by a single physically meaningful B-factor. In contrast, poorly ordered regions would exhibit will exhibit substantially higher RMSD values and physically meaningless B-factors as a result of the refinement’s inability to converge on a single solution.

### Resolution devolution: generating lower resolution structures in silico from near-atomic cryo-EM data

In order to establish a quantitative baseline correlation between the precision of an atomic ensemble and the resolution of an EM density, we first generated a standardized set of EM structures using previously published and universally available datasets. It is expected that, without implementation of external molecular dynamics (MD) force fields, modeling and refinement of structures into maps at 5 Å resolution or worse is ill advised (Goh 2016). Cryo-EM maps at lower than 5 Å resolution have been the targets of a multitude of MD-centric refinement packages, and numerous groups have recently proposed the implementation of MD methodologies to validate the accuracy of atomic models generated from EM maps (Joseph 2016, Singharoy 2016). However, since the majority of near-atomic resolution structures are being build de novo from the cryo-EM density using software such as COOT (Emsley 2010) and refined using crystallographic packages that have been modified to work with cryo-EM densities (Adams 2010, Brown 2015, Wang 2016), we set a lower boundary of 5 Å for the generation of our set of EM densities and convergence tests.

Electron Microscopy Public Image Archive (EMPIAR) (Iudin 2016) datasets 10025 (20S proteasome) (Campbell 2015) and 10061 (β-galactosidase) (Bartesaghi 2015) were downloaded, preprocessed using the Appion processing environment (Lander 2009), and refined using RELION (Scheres 2012) to ~2.7 Å and ~2.2 Å resolution, respectively (see **Methods, Figure 1, Figure 1– figure supplement 1, and Figure 1–figure supplement 2**). To generate lower-resolution structures from these datasets, a random translational offset within a defined range was applied to the refined particle coordinates (see **Methods**). The degree of random translational offset was adjusted empirically for each dataset to generate numerous maps at a range of resolutions. Using this methodology, 14 structures of the 20S proteasome from ~2.7 Å to ~4.9 Å resolution and 14 structures of β-galactosidase from ~2.2 Å to ~4.9 Å resolution were generated.

**Figure 1.**
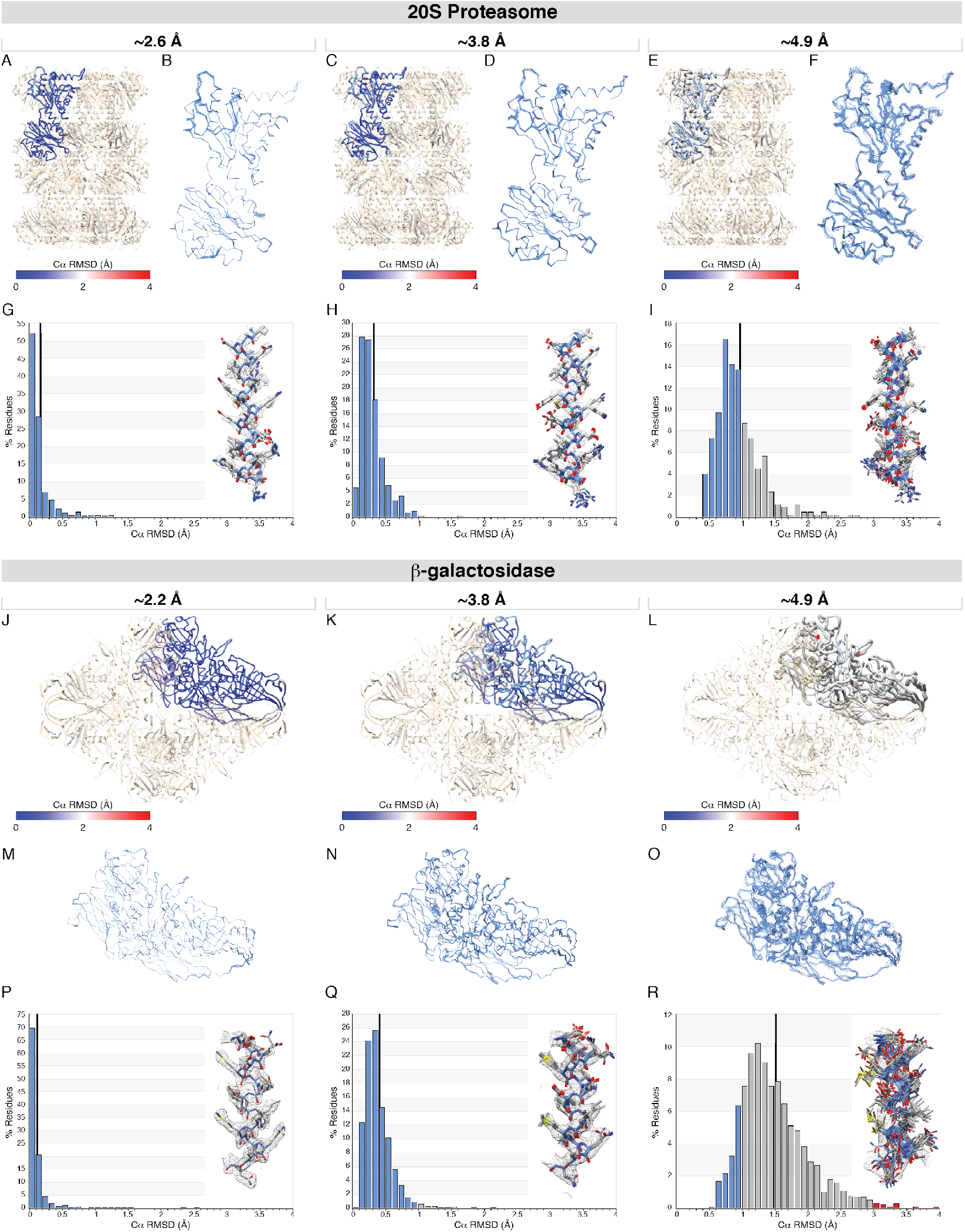
20S proteasome and β-Galactosidase RMSD (C_α_) analysis across various resolutions. The results of the multi-model pipeline and corresponding RMSD (C_α_) analysis are presented for the 20S proteasome core and β-galactosidase, each at three different resolutions –20S proteasome at ~2.7 Å resolution (A, B, and G), at ~3.3 Å resolution (C, D, and H), at ~4.9 Å resolution (E, F, and I), and β-galactosidase at ~2.2 Å resolution (J, M, and P), at ~3.3 Å resolution (K, N, and Q), at ~4.9 Å resolution (L, O, and R). For each structure, the ASU is shown in worm representation and colored by per-residue RMSD value (panels A, C, E, J, K, and L). Line representation for the backbone atoms for the top 10 models of each ASU for each EM density are shown (panels B, D, F, M, N, and O). For each structure, a per-residue RMSD (C_α_) histogram plot is displayed (panels G, H, I, P, Q, and R) where RMSDs less 1 Å are shown in blue, RMSDs between 1-3 Å are shown in gray, and RMSDs greater than 3 Å are shown in red. The mean per-residue C_α_ RMSD value is shown as a black vertical bar. Panels G, H, and I *insets*: residues 49-72 of the 20S proteasome β-subunit are shown in stick representation with the corresponding EM density (zoned 2 Å around the models) shown as a gray mesh. Panels P, Q, and R *insets*: residues 368-383 of β-galactosidase are shown in stick representation with the corresponding EM density (zone 2 Å around the models) shown as a gray mesh. For all stick representations, the backbone atoms are colored in blue with the side chain atoms shown in dark gray.

A comparison of these “simulated” lower resolution maps with previously published reconstructions at corresponding resolutions, i.e. 20S proteasome structures at ~3.3 Å and ~4.8 Å resolution (EMD-5623 (Li 2013) and EMD-6219 (Wang 2015), respectively) and β-galactosidase at ~3.2 Å resolution (EMD-5995 (Bartesaghi 2014)), show comparable quality of density and FSC curves (**Figure 1–figure supplement 3 and Figure1–figure supplement 4**). The asymmetric unit (ASU) from each of the generated lower resolution 20S proteasome and β-galactosidase structures are shown in **Figure 1–figure supplement 5 and Figure1–figure supplement 6**, respectively. Comparison of these densities shows a gradual but noticeable decline in resolvable side chain and backbone features, as anticipated (**Figure 1–figure supplement 5 and Figure1–figure supplement 6**). Together, these comparisons corroborate the quality of the multi-resolution suite of maps generated in this study.

### User-independent pipeline to generate multiple atomic models using cryo-EM maps

This array of EM densities served as a testing suite to quantify the influence that map resolution has on convergence of atomic models. Initial models for refinement using Rosetta were generated by stripping the associated PDB files (PDB IDs: 1YAR (Forster 2005) and 5A1A (Bartesaghi 2015) for 20S proteasome and β-galactosidase, respectively) of cofactors (i.e. PA26, non-protein atoms and ligands, etc.), removing alternate conformations, setting all occupancies to one, resetting all B-factors, and correcting Ramachandran and geometric outliers. These initial models were then refined into the EM density using the corresponding symmetry and resolution values while adjusting the Rosetta weighting and scoring functions according to the estimated map resolution. In order to sample a large conformational space for each given map, regardless of resolution, 100 models were generated using Rosetta (Figure 2). A comparison of the top 10 structures resulting from 100 or 1000 Rosetta-generated models using ~2.8 Å and ~4.8 Å resolution densities were essentially indistinguishable (see **Figure1–figure supplement 7**), indicating that 100 models are sufficient for these convergence analyses.

**Figure 2.**
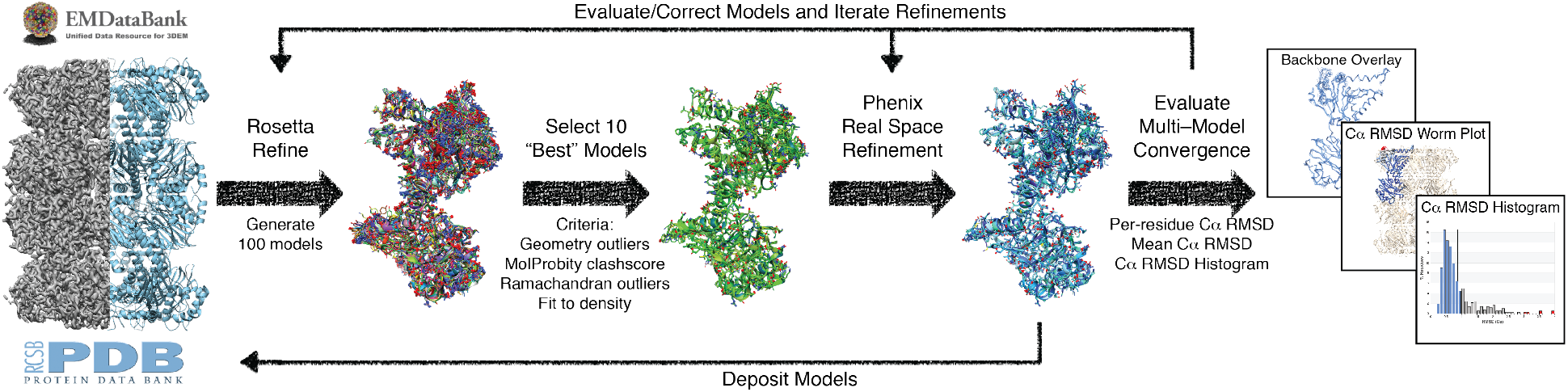
Overview of the multi-model “pipeline”. For each EMD entry, the corresponding PDB entry was stripped of non-protein atoms and alternate conformers followed by resetting all occupancies to 1 and B-factors to the approximate mean value. This initial model was then refined against the primary EMD map entry using Rosetta with a desired output of 100 models. The statistically “best” ten models – the 10 structures that had the fewest geometric outliers, fewest Ramachandran outliers, the lowest MolProbity clashscore, and the best Rosetta aggregate score – were then subjected to real-space refinement in Phenix. The per-residues RMSD (C_α_) was then calculated for each model against the refined structures. Thick arrows indicate the linear pipeline employed in this study while the thin arrows represent the iterative process that could be utilized by a general user to generate a previously unpublished structure.

The 100 Rosetta-generated models were ranked based on the number of Ramachandran outliers (%, lower better), geometry violations (%, lower better), Rosetta aggregate score (value, lower better), and MolProbity clashscore (Chen 2010) (value, lower better). These criteria were chosen in an effort to select models that possessed good geometry as well as good model-to-map agreement. The 10 structures that scored the best across all categories were selected for real-space refinement using Phenix (Adams 2010). Real-space refinement of coordinates and B-factors was performed because the resulting structures from Phenix generally yielded better geometries and model-map correlation coefficients than the Rosetta models. However, algorithm and parameterization improvements in recent and future releases of Rosetta may remove the need for this additional refinement step (Wang 2016).

Following real-space refinement using the Phenix suite, the per-residue C_α_ root-mean-square deviation (RMSD) was calculated for all residues in the asymmetric unit of the refined structures (**Figure 1 and Figure 2**). The all-atom RMSD values were also calculated, with these values typically greater than C_α_ RMSD values by ~1-1.5 Å (data not shown). However, because residues comprise different numbers of atoms and some protein side chains possess internal symmetry (i.e. phenylalanine or tyrosine) resulting in ambiguous atom correspondence, C_α_ RMSD was used for these analyses. In total, 10 models were used to derive validation statistics for all densities in this testing suite, providing a measure of unity in atomistic positions within the asymmetric unit relative to reported resolution. To provide a general overview of the atomic convergence of each structure, the mean C_α_ RMSD value and a C_α_ RMSD histogram were also calculated for each map (**Figure 1, Figure 1–figure supplement 8, and Figure 1–figure supplement 9**).

### Multi-model agreement provides global and per-residue metrics for assessing cryo-EM map quality

Comparison of the multi-model agreement statistics for each set of atomic models generated against the lower resolution structure suite provides insights into how these global and per-residue model quality metrics can inform on the precision of single atomic coordinates for structures refined against EM densities spanning a broad resolution range (**Figure 1**). As anticipated, the multi-model agreement statistics directly correlate with map resolution, with the mean C_α_ RMSD of each group of models increasing as the resolution of the map decreases from ~2.2 Å to ~4.9 Å (**Figure 1, Figure 1– figure supplement 8, and Figure 1–figure supplement 9**). For example, the mean C_α_ RMSD value increases gradually from 0.30 Å for the ~2.7 Å resolution 20S proteasome dataset, to 0.36 Å and 1.14 Å for the ~3.8 and ~4.9 Å resolution datasets, respectively (**Figure 1G, H, I, and Figure 1–figure supplement 8**). A similar trend is also observed for the β-galactosidase datasets, with the mean C_α_ RMSD increasing from 0.25 Å to 1.76 Å as the resolution of the map decreases from ~2.2 Å resolution to ~4.9 Å resolution, respectively (**Figure 1P, Q, R, and Figure 1–figure supplement 9**).

This observed correlation between map quality and model convergence is not unexpected, given that lower resolution maps possess fewer well-resolved features for accurate placement of side chain atoms and, in some cases, are completely absent of side chain densities (**Figure 1–figure supplement 5 and Figure 1–figure supplement 6**). Furthermore, EM maps reported to worse than ~5 Å resolution often contain secondary structure elements that are nearly feature-less, leading to ambiguous β-strand or α-helix register (Hryc 2011). In agreement with these observations, comparison of the same α-helix from the “best” 10 models and corresponding EM density from each of the 20S proteasome and β-galactosidase structures reveals decreasing multi-model agreement as the resolution of the map worsens (**Figure 1**). Specifically, the pronounced side chain densities that gives rise to near perfect multi-model agreement for the highest resolution structures (**Figure 1G, P**) are diminished in the lower resolution structures (**Figure 1I, R**), corresponding to increased heterogeneity in backbone and side chain atom placements (i.e. increased mean and per-residue C_α_ RMSDs), correlated with worsening map resolution.

Although EM densities are ascribed a single resolution value, the local resolution is never isotropic across the entire map, with most structures exhibiting a marked range of varying local resolution (Leschziner and Nogales 2007, Adams 2010). Even for a protein such as the 20S proteasome from the thermophilic bacterium *Thermoplasma acidophilum*, which possesses high internal symmetry (D7) and high thermal stability (thermoinactivation ~97 ˚C) (Beadell and Clark 2001), there is nonetheless a range of local resolution and thus varying map quality (**Figure 1–figure supplement 8 and Figure 3**). Due to the direct correlation between multi-model agreement and EM density quality (*vida supra*), we reasoned that examination of the multi-model agreement statistics at a per-residue level would inform on the local map quality. In support of this notion, the per-residue C_α_ RMSD values calculated from the multi-model refinements were mapped to the ASU (**Figure 1 A, C, E, J, K, and L**). Each structure is represented using a tri-color heat map coloring scheme with residues exhibiting C_α_ RMSDs less then 1 Å colored blue, between 1-3 Å colored gray, and greater than 3 Å colored red. For both the 20S proteasome and β-galactosidase datasets, regardless of the overall resolution estimate, the regions exhibiting the most heterogeneity (highest per-residue C_α_ RMSD values) are localized to the regions least well resolved, as evidenced by the poorer EM density and lower local resolution estimates (**Figure 1, Figure 3, Figure 1–figure supplement 8, and Figure 1–figure supplement 9**).

**Figure 3.**
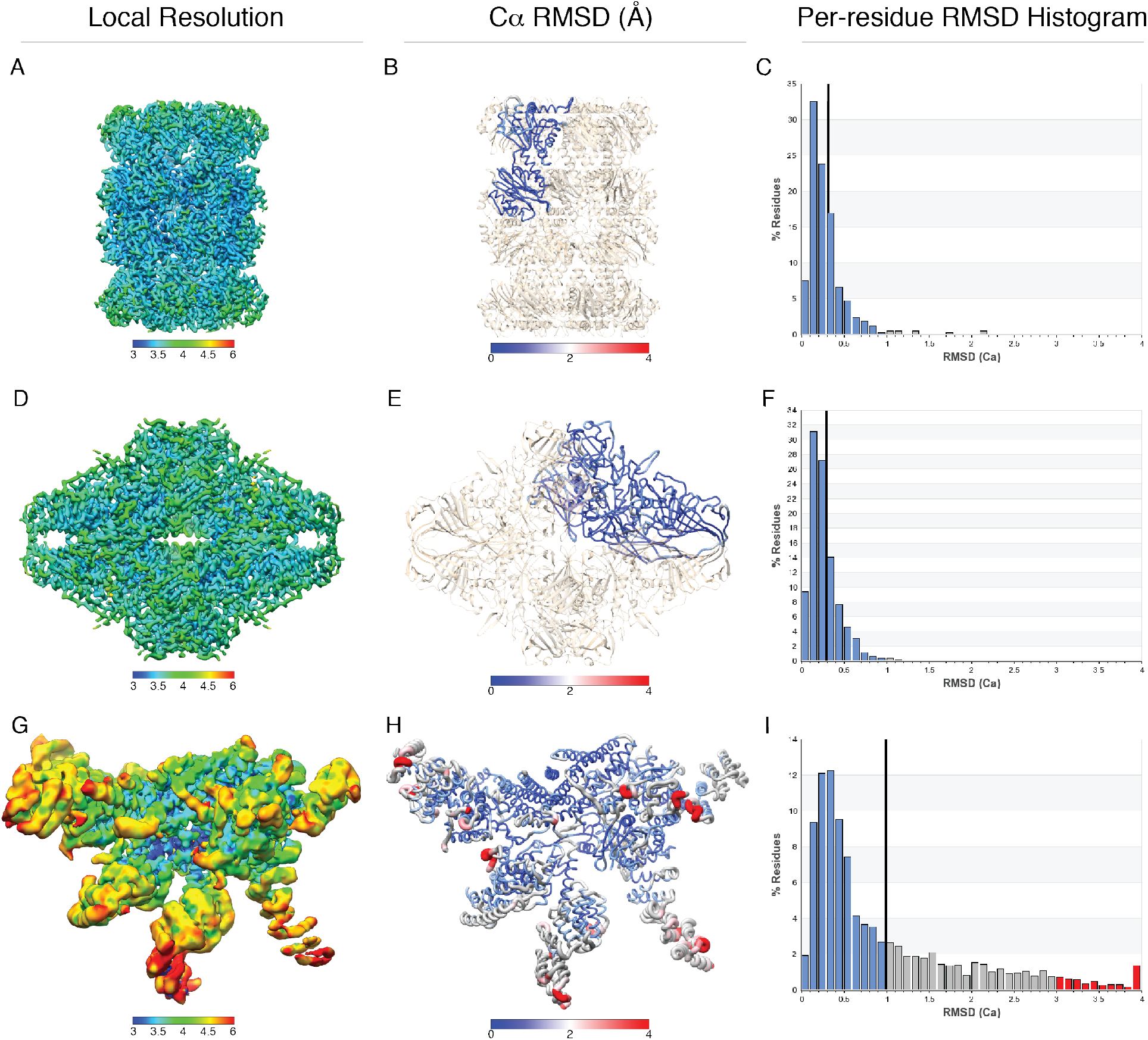
Influence of map resolution variance on convergence of atomic models. 3D local resolution plots for cryoEM maps of 20S proteasome (panel A), β-galactosidase (panel D), and EMD entry 6479 (panel G) having overall FSC-reported resolutions of ~3.5 Å. A worm plot and histogram of the per-residue C_α_ RMSDs, (panels B, E, and H and panels C, F, and I, respectively) are shown for each map. The worm plots show that areas exhibiting increased per-residue C_α_ RMSDs are correlated with lower local resolution estimates, as expected. The histograms show that as the range of resolutions contained within a map increases, the right-skewness of the histogram also increases along with the separation of the mean (black line) and mode of the histogram.

The C_α_ RMSD histogram provides an easily interpretable means to quickly evaluate both the overall quality of an EM map, as well as informing on the range of resolutions contained within the map. An overall shift of C_α_ RMSD histogram to the right, where the mean and mode of the RMSD values are similar, is indicative of a general worsening in map resolution, exemplified by an increasing number of residues possessing greater than 1 Å C_α_ RMSD values (**Figure 1G, H, I, P, Q, R, Figure 1–figure supplement 8, and Figure 1–figure supplement 9**). However, a pronounced right-skewness of the per-residue C_α_ RMSD histogram, where the mean and mode diverge, is indicative of a map that contains a wide range of local resolutions (**Figure 3**). Examination of EMD-6479 (Dambacher 2016) corroborates the observation that per-residue C_α_ RMSD values inform on local map quality. Specifically, EMD-6479 exhibits a pronounced decrease in local resolution at the periphery of the molecule, with the core resolved to ~3.2 Å resolution while the outer, peripheral regions are resolved to ~5 Å resolution or worse. Accordingly, the gradual decrease in local resolution as densities extend toward the periphery of the molecule is consistent with a gradual increase in the per-residue C_α_ RMSD values, and a substantial right-skewness of the C_α_ RMSD histogram (**Figure 3**). Together, these analyses indicate that both the mean C_α_ RMSD and profile of the C_α_ RMSD histogram provide a means to globally assess the multi-model agreement and, presumably, overall map quality, with per-residue C_α_ RMSD values reporting on local map quality.

Comparison of the per-residue C_α_ RMSD values and the B-factors assigned from Phenix real-space refinement shows a correlated trend of values, with the highest values reported at peripheral and flexible regions (**Figure 3 – figure supplement 1**). However, the B-factors exhibit a wider range of variability than the C_α_ RMSD values when comparing one structure to another, complicating comparisons of density quality between depositions. Furthermore, estimation of B-factor is not consistent from one refinement package to another, and refinement strategies and parameters can influence the B-factor values even when using a single package. Regardless of the package used, the implied RMSDs for assigned B-factors are generally lower than the RMSDs determined by our analyses. B-factor values considered to be extremely “high” by crystallographic standards (300 to 400), imply an RMSD of ~2 Å (Kuzmanic 2014), which, as demonstrated by the C_α_ RMSD histograms (**Figure 3I**), is an overestimation of the electron density quality present in cryo-EM structures. An RMSD of 3 Å would correspond to a B-factor of over 700, which is not physically reasonable, reflecting the inappropriateness of the isotropic approximation of microscopic heterogeneity. Additionally, potential corrections to scattering factors and a limited understanding of radiation damage further complicate the interpretation of B-factors in cryo-EM densities relative to X-ray crystallography. In contrast, assignment of a per-residue C_α_ RMSD provides a consistent and interpretable reporter on the quality of electron density.

### Evaluation of the EMDB using the user-independent multi-model pipeline

In order to assess the range of map and model quality extant in the cryo-EM field, the multi-model pipeline outlined above was applied to all Electron Microscopy Data Bank (EMDB, www.emdatabank.org) depositions having a reported resolution of better than 5 Å with an associated PDB entry (see **Methods** and **Figure 2**). Analyses were limited to structures exhibiting C- or D-symmetry (i.e. C1, C2, D7, etc., not helical or icosahedral) that did not contain nucleic acids (i.e. ribosomes or spliceosomes, etc.). These structure types were excluded because, at the time this study was initiated, refinement of these structure types using the employed software (Rosetta and PHENIX) were not robust. Current releases of these software packages now accommodate these structure types, so that the implementation and analysis described below is more broadly applicable (Chowdhury 2017).

A scatter plot of the multi-model agreement statistics (mean C_α_ RMSD) for all 139 structures analyzed in this study against their reported resolution value shows a strong correlation, with higher mean C_α_ RMSD values associated with lower resolution structures (**Figure 4**). These observations were anticipated because, as discussed above, the multi-resolution suite of 20S and β-galactosidase structures exhibit a strong resolution–model quality relationship (**Figure 1, Figure 1–figure supplement 8, and Figure 1–figure supplement 9**). Comparison of the mean C_α_ RMSD values determined from the EMDB and *in silico* datasets vs. resolution provide valuable insights into both global and local map quality. Specifically, plotting C_α_ RMSD vs. resolution yields an exponential fit against the data with the apex of the exponential function located at ~4.1Å, for both the EMDB and *in silico* datasets. These observations indicate that beyond ~4.1 Å resolution, where clearly resolvable side-chain density becomes rare, convergence of multiple models against the density decreases precipitously (**Figure 4**). The observation that both the 20S and β-galactosidase datasets, while exhibiting the same general trend as the cumulative EMDB entries, consistently present a ~1.5 Å lower overall RMSD, is both a result of the limited structural heterogeneity/flexibility in the maps and the high quality of the initial atomic model. Specifically, for both the 20S and β-galactosidase datasets, the high resolution X-ray model was refined into the highest resolution EM density (20S PDB ID: 1YAR (Forster 2005), β-galactosidase PDB ID: 1DP0 (Juers 2000)). As a result, the structures are initialized at a lower global minimum than what would typically be obtained for most models built *de novo* into cryo-EM densities at 3.5 Å resolution or worse.

**Figure 4.**
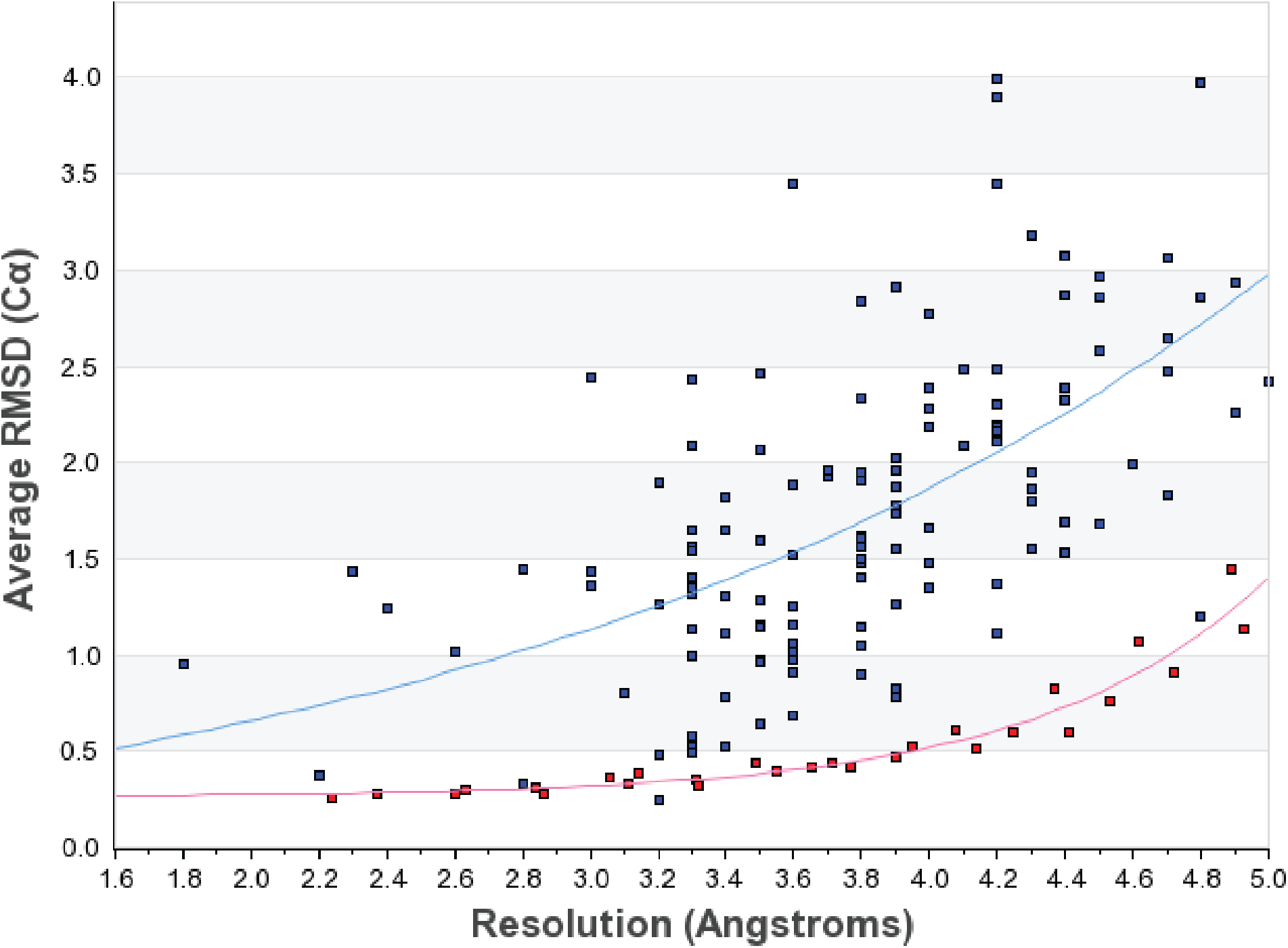
Average RMSD (C_α_) versus FSC-reported resolution. The average RMSD (C_α_) value calculated from the “best” 10 models resulting from the multi-model pipeline plotted versus the FSC-reported resolution of the target map. Values for EMDB entries are shown as blue squares and values for the *in silico* datasets are shown as red squares. The red line is a non-linear regression fit against the 20S and β-galactosidase structures generated in this study. The cyan line is a non-linear regression fit against the EMDB entries.

A non-linear regression fit of mean C_α_ RMSD values versus resolution provides an estimated mean C_α_ RMSD value for models refined against an EM density for various reported resolutions between ~2 – 5 Å resolution (**Figure 4**). As a result, structures giving rise to mean C_α_ RMSD values that lie substantially above the regression line are likely to possess EM density that is worse than anticipated, given the FSC-reported resolution. Whether the density is poorer than expected for the given resolution, or if the reconstruction contains large local resolution variations, can be assessed by examining the per-residue RMSDs and RMSD histogram for the entry (**Figure 1, Figure 3, Figure 1– figure supplement 8, and Figure 1–figure supplement 9**). These elevated RMSD values may also be due to modeling errors introduced by the user-independent, automated pipeline.

It is apparent from this plot that most structures (72 %) worse than ~4.5 Å resolution have a mean C_α_ RMSD value of 2.0 Å or above by our analyses. Such RMSD values arise from significant backbone displacements between models, often with little agreement in side-chain placement. Regardless of resolution, maps giving rise to a mean C_α_ RMSD of greater than 3 Å show significant heterogeneity in backbone atomic coordinates, with almost no agreement in side-chain placement for a majority of the map. Great caution and consideration should be exercised in the interpretation of such models. In contrast, some maps with a reported resolution of worse than 4.0 Å resolution yielded mean C_α_ RMSDs that were lower than 2 Å (EMD-8188, EMD-3180), suggesting an adequate degree of confidence in the agreement between models and thus, the approximate placement of atoms in much of the map.

Supporting the reliability of our *in silico* reconstructions and the resulting non-linear regression fit, analysis of the outputs from the multi-model pipeline reveal that the C_α_ RMSD values for 20S proteasome and β-galactosidase EMDB entries (EMD-2984, -5623, -5995, -6219, and -6287) lie on the same trend line as the *in silico* datasets. Maps estimated to have similar resolutions (i.e. the deposited ~4.8 Å resolution 20S proteasome EMDB entry 6219 and the ~4.7 and ~4.9 resolution *in silico* 20S proteasome maps) report C_α_ RMSD values within the standard error of the measurements (**Figure 4**). These comparisons not only verify that multi-model analyses can report on map quality but also corroborate the ability of the *in silico* suite of structures to recapitulate the features of cryoEM structures at a range of resolutions.

### Not all maps are created equally – Case studies at 2.8 Å and 4.8 Å resolution

In an effort to report on the variations in the quality of EM density for maps that are reported to have identical resolutions, as well as to test the robustness of this methodology at a range of resolutions, we focused our multi-model agreement analyses on EMDB entries reported at ~2.8 Å and ~4.8 Å resolution (**Figure 5**).

**Figure 5.**
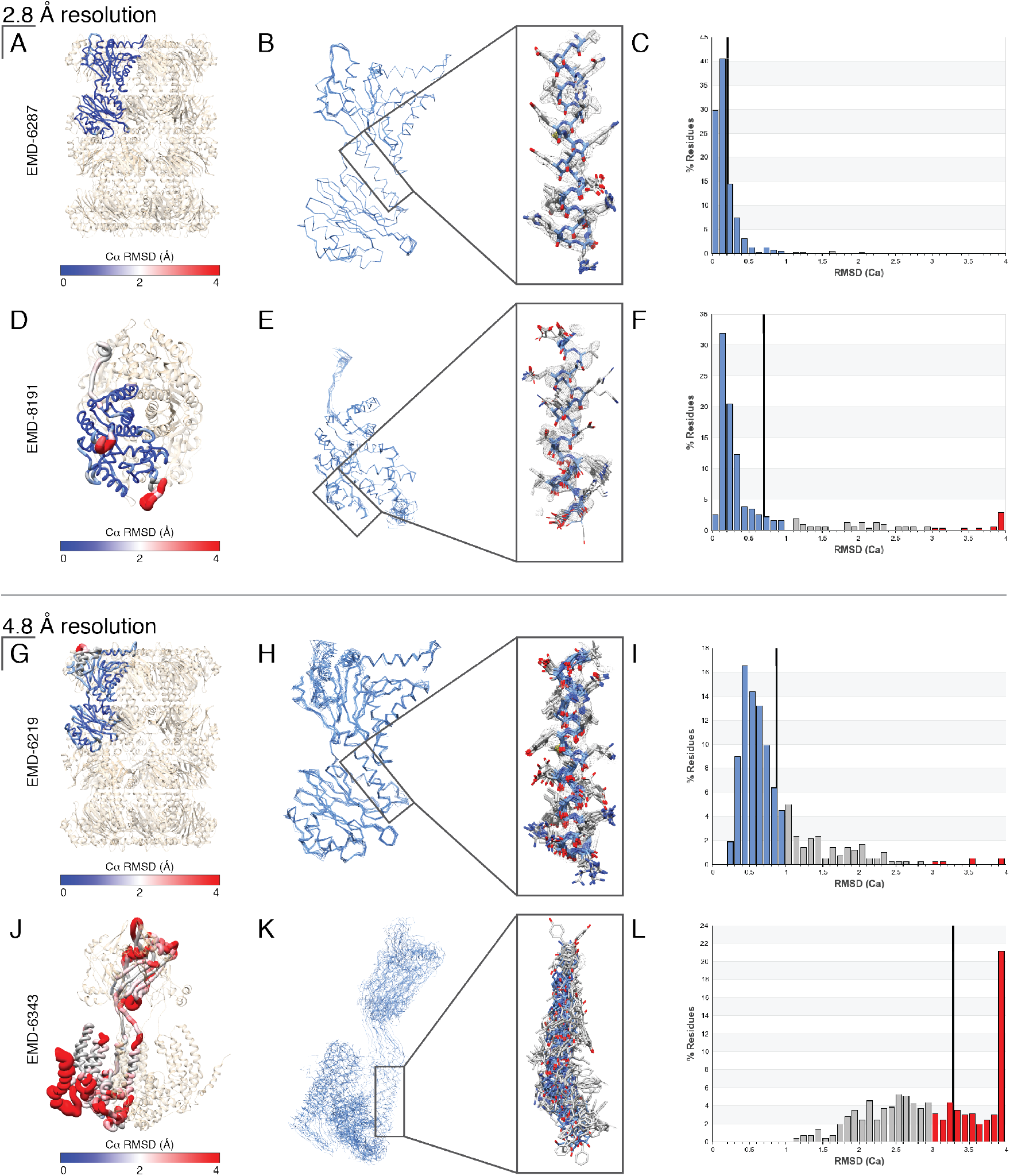
Variation in map quality at similar FSC-reported resolution values – Comparisons of maps at 2.8 Å and 4.8 Å resolution. Comparison of the multi-model analysis for structures at 2.8 Å resolution – EMD entries 6287 (panels A-C) and 8191 (panels D-F) – and at 4.8 Å resolution – EMD entries 6219 (panels G-I) and 6343 (panels J-L). For each structure: (Left panel) the ASU is shown in ribbon representation with each residue colored by the per-residue RMSD (C_α_) value (the rest of the molecule is colored in wheat), (Middle panels) the ASU from the top 10 models are shown in line representation with a single α-helix (inset, helix atoms exhibit approximately the mean C_α_ value for the entire molecule) and corresponding EM density (zoned 2 Å around atoms) shown in gray, and (Right panels) the per-residue RMSD (C_α_) histogram plot. For all stick representations, backbone atoms are colored blue with side chain atoms shown in gray. For the histograms, the average per-residue C_α_ RMSD value is shown as a black vertical bar.

EMDB entries 6287 (Campbell 2015) and 8191 (Merk 2016) were both reportedly determined to ~2.8 Å by FSC. Our analyses, as expected, yielded a suite of models for these two entries with mean C_α_ RMSD values < 1 Å (0.33 and 0.69 Å, respectively). EMDB entry 8191 shows a larger variation in local map quality, with several loops and secondary structure elements at the periphery more poorly resolved than the core of the molecule. As a result, the per-residue C_α_ RMSD values in these regions are elevated above the mean, as evidenced by: more gray and red regions when coloring each residue by the per-residue C_α_ RMSD value (**Figure 5D**); by increased heterogeneity in the atomic coordinates across the “best” models (**Figure 5E**); and by the right-skewed histogram (**Figure 5F**). In contrast, EMDB entry 6287 shows little variation in local map quality and, as a result, yields a narrow RMSD histogram with only ~3% of residues with C_α_ RMSDs greater than 1 Å (**Figure 5A, C**). By comparison, EMDB entry 8191 yields ~10% of residues with per-residue C_α_ RMSD values greater than 1 Å (**Figure 5F**). In addition, comparison of extracted regions from both EM maps (zoned 2 Å around an α-helix with an approximate mean C_α_ RMSD as the entire molecule) indicate that EMDB entry 8191 has a lower percentage of well resolved side chain densities, as well as broader backbone density as compared to EMDB entry 6287 (**Figure 5B, E**).

The variations in map quality that give rise to large discrepancies in multi-model agreement become much more pronounced for structures reported at resolutions worse than ~4 Å. An example of this is demonstrated by our analyses of EMDB entries (Wang 2015) and 6343 (Ge 2015), both at a reported resolution of ~4.8 Å. Analysis of EMDB entry 6219 shows high multi-model agreement for most of the ASU, with a majority of the ASU possessing per-residue C_α_ RMSDs less than 1 Å (**Figure 5I**). There are several poorly resolved densities at the periphery of EMD-6219, which give rise to elevated RMSDs, shown in red (**Figure 5G, I**). However, a majority of the map possesses well-resolved backbone density, allowing for consistent placement of backbone atoms (**Figure 5H**). In contrast, analysis of EMDB entry 6343, which is also reportedly resolved to ~4.8 Å resolution, shows predominantly elevated per-residue C_α_ RMSDs, with a mean C_α_ RMSD value of 3.9 Å, with no residues better than 1 Å C_α_ RMSD. Over 48% of the residues yielded C_α_ RMSD values greater than 3 Å (**Figure 5J, L**). Examination of the EM density around an α-helix (with an approximate mean C_α_ RMSD of the entire molecule, middle panel) located adjacent to the symmetry axis contains little-to-no discernable side chain density (**Figure 5K**). As a result, the backbone atoms vary significantly across the models, with some helices distant from the three-fold symmetry axis showing significant translation of the α-helix backbone and/or complete changes in helical register.

Together, these data show the benefits of using multiple models to assess the quality of EM maps, presenting a resolution-independent and clearly interpretable visualization of how well resolved the density used to construct a model is, as well as providing a level of confidence for the assigned positions of side chains.

### Examination of EMDB entry 3295

The 20S proteasome and β-galactosidase structures detailed above are highly symmetric and exhibit less variation in local resolution than most entries in the EMDB (**Figure 3, Figure 1–figure supplement 8, and Figure 1–figure supplement 9**). EMDB entry 3295 exemplifies a deposition reported to be at high resolution with an associated full atomic model (PDB ID: 5FTJ), but which also exhibits a wide range of local resolution in the map (Banerjee 2016). The overall resolution of the map is reported to be 2.3 Å, however, unlike the 20S proteasome or β-galactosidase, the quality of the map exhibits extremely internal variability, with external loops and an entire peripheral domain that are poorly resolved (**Figure 6**).

**Figure 6.**
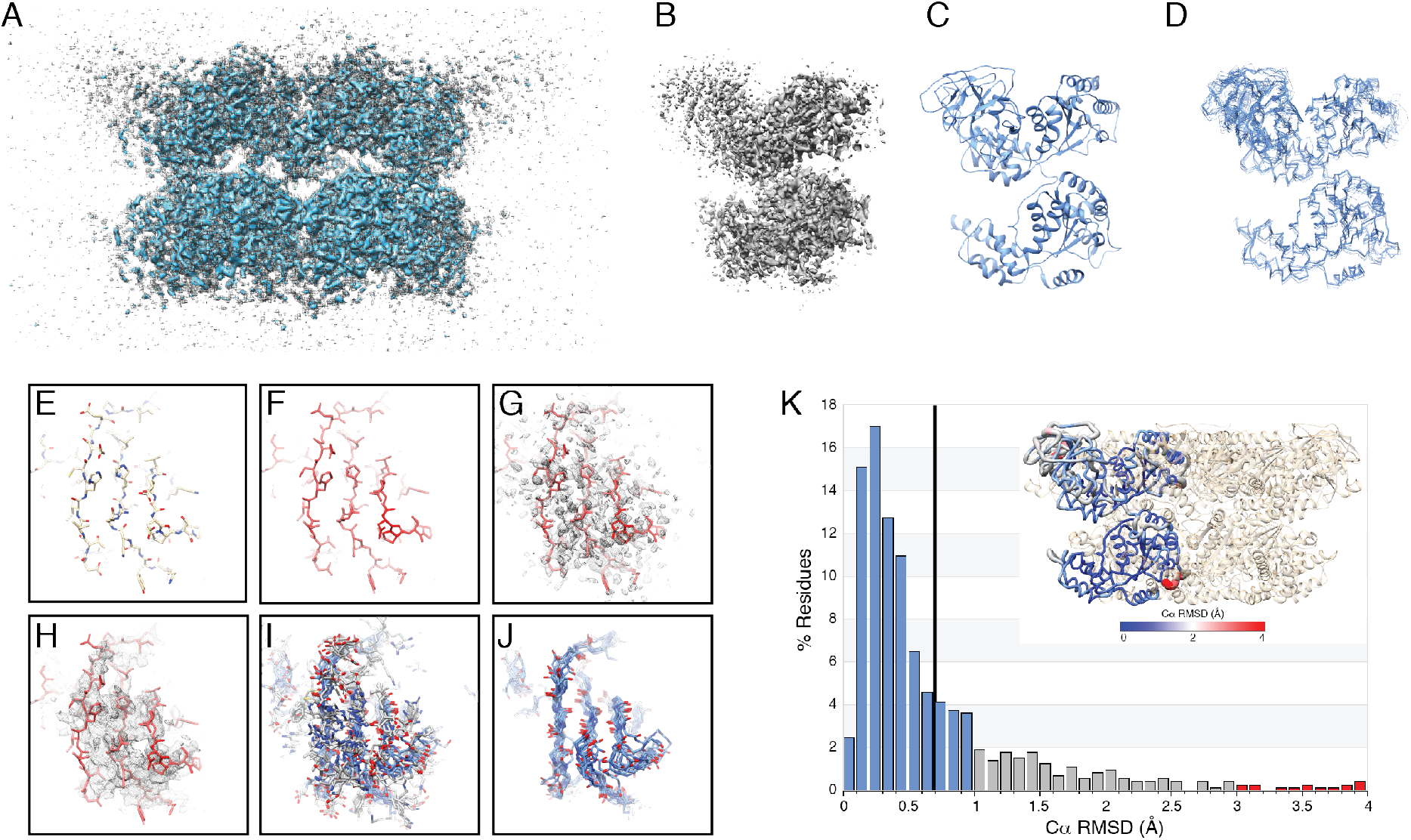
Case study: EMD-3295. (A) The primary map for entry EMD-3295 displayed at two different contour levels –lower threshold (gray) and higher threshold (blue) – showing loss of density for the N-terminal domain upon increasing threshold. (B) Corresponding EM density from EMD-3295 for a single ASU (zoned 5 Å around chain A). (C) Chain A from PDB entry 5FTJ shown in ribbon representation. (D) Backbone atoms of chain A from the top 10 models generated by the multi-model pipeline shown in line representation. The same region of PDB entry 5FTJ shown in stick representation (E), colored by C_α_ B-factor (F), with the sharpened EM density from EMD-3295 shown as gray mesh (G), with the unsharpened EM density from EMD-3295 shown as gray mesh (H), the top 10 models from this study shown in stick representation (I), and only backbone atoms (J). For panels (I) and (J), the backbone atoms are colored blue with the side chain atoms shown in dark gray. (K) Histogram plot of the per-residue RMSDs (C_α_, RMSDs less 1 Å shown in blue, RMSDs between 1-3 Å shown in gray, and RMSDs greater than 3 Å shown in red) with the ASU shown in ribbon representation and colored by per-residue RMSD (K, inset). For the histogram, the average per-residue C_α_ RMSD value is shown as a black vertical bar.

EMD-3295 is a homohexamer containing a well-ordered hexamerization domain, as well as an N-terminal domain that extends from the core that is resolved to a much worse resolution than the reported ~2.3 Å resolution (**Figure 6A**). The disorder of this domain is clearly evident when one extracts the EM density within 2 Å radius of the atomic model of a single ASU (**Figure 6B**), shown as a cartoon in **Figure 6C**. The EM density and associated atomic model were subjected to the linear multi-model pipeline described above and, as anticipated, there is excellent multi-model agreement within the core of the protein, and lower levels of agreement in the poorly resolved N-terminal domain (**Figure 6D**). These observations are corroborated by the varied conformational heterogeneity across the resulting multiple models (**Figure 6D**), the per-residue C_α_ RMSD worm plot (**Figure 6K**, inset), and the pronounced right-skewness of the per-residue C_α_ RMSD histogram (**Figure 6K**). Furthermore, comparison of the per-residue C_α_ RMSDs for this domain to the rest of the molecule (**Figure 6K**) shows that most of the disordered N-terminal domain have C_α_ RMSD values above the mean, with the highest C_α_ RMSD values localized almost entirely to the N-terminal domain and two peripheral loops.

The multi-model analysis of EMD-3295 clearly outlines the rationale for depositing multiple atomic models for cryo-EM densities. A ~2.3 Å resolution structure would be considered by the broad structural biology community to be of high quality, even in regions that are assigned high B-factors. However, it would be imprudent to carry out a detailed interpretation of the deposited atomic coordinates within the N-terminal domain due to the extremely poor density in this domain. A naïve user could examine the panels in **Figure 6** and rapidly deduce that the N-terminal region of this map is poorly ordered, and consider the low confidence of atom positioning when interpreting the model. Furthermore, if the same naïve user is presented with 10 or more atomic models upon downloading the PDB entry associated with cryo-EM map for the purposes of, for example, designing a mutagenesis experiment, the user would intuitively be able to distinguish between alternative interpretations embodied by the widely divergent models of poorly ordered regions and the converged high-confidence models for well-ordered regions.

## Discussion

Resolution estimates for EM maps refined using the gold-standard FSC refinement protocol (Henderson 2012, Scheres and Chen 2012) can provide an overall estimate of the quality of a map, but it is important to note that this measure only reports on the self-consistency between the two independent half reconstructions obtained during refinement. Thus, EM maps suffering from pathologies such as anisotropic resolution (i.e. gaps in Euler space) or large local resolution variations will report inflated resolution estimates with markedly lower resolution along one or more axes or contain entire domains that are very poorly resolved, respectively (Penczek 2002, Lander 2013, Banerjee 2016). In the case of the latter, the resolution estimate for a map can further be inflated by applying a 3D mask to exclude poorly resolved regions of the map from the FSC calculations. As such, ascribing a singular value to an EM density as the sole means to inform on its quality is inadequate, and other metrics are needed to assess both EM map and model quality.

Here, we present a straightforward methodology using readily accessible software packages to generate multiple models, each of which represent an approximately equally valid interpretation of the EM density. The principle conclusion from our work is that examination of the convergence of multiple structures provides a means to assess both global and local EM map quality that is independent of FSC. Specifically, as the resolution of the target map worsens and the extent of resolvable side chain densities decreases, the agreement in atomic coordinates across the multiple models decreases accordingly, with lower resolution structures exhibiting higher mean and per-residue C_α_ RMSDs (**Figure 1, Figure 1–figure supplement 8**, and **Figure 1–figure supplement 9**). Importantly, this correlation is not limited solely to global map quality, i.e. nominal FSC-reported resolution, as the per-residue C_α_ RMSD values correlate well to local resolution estimates. As such, maps exhibiting large variations in local resolution also exhibit large variances in per-residue C_α_ RMSD values, with the least resolved regions of the maps also exhibiting the highest per-residue C_α_ RMSD values (**Figure 3**).

We expect that an important use of the multiple models generated using these methodologies will be to define the range of structures that are compatible given the EM density. Presumably, each model within the ensemble is of approximately equal quality, with all models exhibiting reasonable geometries and equal fits to the density. Because each model is generated using essentially the same calculation, the differences between the multiple models reflect the reproducibility of the refinement methodologies given the data, and provide a lower limit of the uncertainties of the structure calculation (Terwilliger 2007). As such, the conclusions based on one model from the group of models are as likely to be correct as the conclusions drawn from any other model within the group. However, comparison of the atomic coordinates across the ensemble is more informative and provides a better platform for drawing conclusions, minimizing the risk of structural over-interpretation resulting from the seemingly rigid single-structure models.

Visual examination of multiple models provides an immediate qualitative metric to even the most naïve user. Specifically, the general user typically evaluates the “correctness” of an atomic model using a variety of metrics, including global model statistics (i.e. MolProbity score (Chen 2010), map correlation coefficient, map-to-model FSC (DiMaio 2013), Ramachandran outliers, EMRinger score (Barad 2015), etc.), in addition to local examination of the model using B-factor analysis and fit to density. However, as described through our examination of EMD-3295, evaluating the “correctness” of an atomic model using B-factor analysis can be misleading about the extent of conformational heterogeneity present in the map. In the absence of the corresponding EM density, conventional models cannot always inform the general user of areas of poor model/map quality and/or poor model–map agreement. In contrast, evaluation of the multiple independent atomic models, even in the absence of the corresponding EM density, immediately provides a qualitative metric as to the “correctness” of the atomic coordinates – where regions or poor map quality will exhibit a large degree of heterogeneity in the atomic coordinates of the ensemble due to an inability of the refinement to converge to a single solution.

A need for this type of intuitive and accessible reporter on intrinsic structural heterogeneity is made apparent through our analyses of the deposited models in the EMDB. The multi-model convergence criterion for structures refined against the *in silico* 20S proteasome and β-galactosidase suite of EM densities show a strong correlation with map quality (i.e. global resolution), due in part to the fact that these complexes do not contain regions of substantial flexibility. The macromolecular structures deposited to the EMDB show a much higher degree of variability in the relationship between model convergence and reported global resolution. While non-linear regression analysis of *in silico* datasets and the previously deposited EMDB entries possess the same shape exhibited by the *in silico* datasets, the mean C_α_ RMSD values for the EMDB entries are, on average, ~1 Å higher than the *in silico* datasets. This is not entirely unexpected, as the *in silico* datasets represent a best-case modeling scenario, with very stable and isotropically resolved maps, and the initial model derived from a high-resolution X-ray crystal structure. However, this disparity between the *in silico* datasets and the EMDB may be lessened with improvements in refinement algorithms, improvement in the accuracy of atomic electron scattering coefficients used during refinement, as well as more robust methods for identification of modeling errors.

Although the methodology we used to generate multiple models in this study followed a linear pipeline, this methodology could be easily modified for all stages of model building and refinement to eventually yield an ensemble of models that best represent the data (**Figure 2**). At the very least, if examination of the per-residue C_α_ RMSD reports regions exhibit greater than 4 Å C_α_ displacements, inclusion of these regions in the final deposited models should be carefully reconsidered or deposited as a C_α_ trace. As evidenced in **Figure 7**, regions of the model exhibiting 3 Å or worse per-residue C_α_ RMSD values can show substantial shifts in amino acid sequence register.

**Figure 7.**
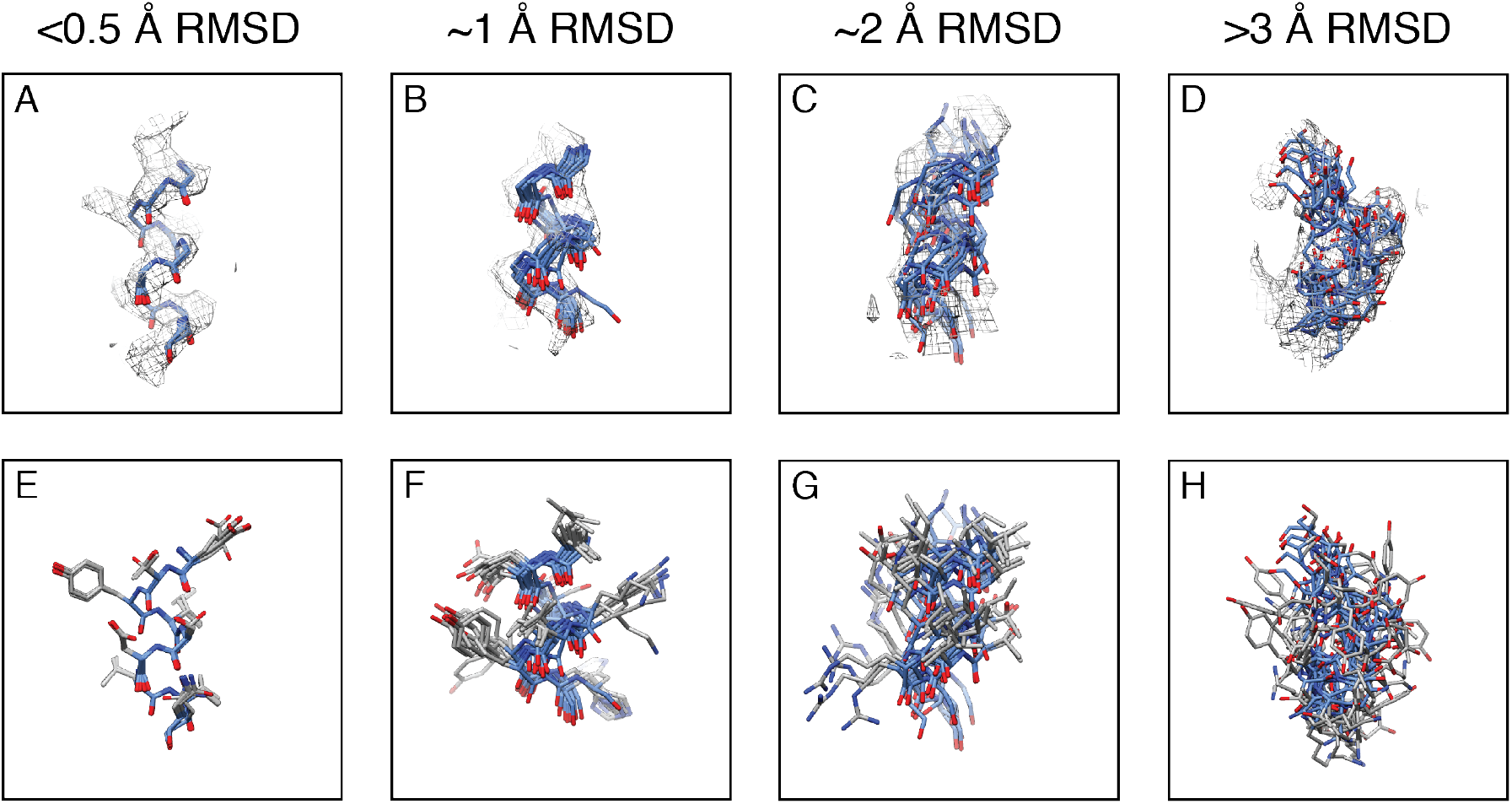
Multi-model convergence for given C_α_ RMSD values. (A-D) For each C_α_ RMSD value, backbone atoms from the top 10 models are displayed with corresponding EM density (zoned ~4 Å around displayed atoms). (E-F) The same residues displayed in panels A-D are displayed with side-chain atoms. For all stick representations, backbone atoms are colored blue with side chain atoms shown in gray.

Current EM data processing and refinement strategies typically strive to determine the most homogenous set of particles and their optimal alignment parameters, but any unresolved conformational and compositional heterogeneity or misalignment of the molecule will be manifested in the final EM density as a decrease in global and/or local resolution. It is important to note that the models generated using these methodologies do not necessarily report directly on the source of the heterogeneity within the data, i.e. dynamics or particle misalignment, and the interpretation thereof should be limited. Rather, the uncertainty of the model coordinates provides a lower bound as to the precision of the models, informing on both map and model quality (DePristo 2004, de Bakker 2006, Ondracek and Mesters 2006, Terwilliger 2007).

These results represent an attempt to depart from the limitations of ascribing a map “quality” metric that is solely dependent on FSC. The *in silico* datasets generated in this study not only provide a means to establish the correlation between EM map quality and model precision, but may serve as test suite for the development of refinement algorithms and alternative metrics to assessing EM map quality that could be used in combination with FSC-reported resolution and established model quality metrics. Nonetheless, the multi-model approach provides an easily accessible mechanism for assessing EM map quality that even the most naïve user can appreciate. Importantly, as has been discussed previously for ensembles refined against X-ray diffraction data, the PDB is amenable to deposition of multiple models and has tools in place for handling and annotating multi-model entries (response associated with (Furnham 2006)).

Beyond this study, we postulate several important developments to the multi-model approach will make this analysis more robust. These include, but are not limited to: methodologies to apply these analyses to datasets that benefit from the combination of multiple EM maps (i.e. focused classification and/or refinements, regions that benefit from different sharpening values, etc.), or methodologies to assess which minimal combination of the best-scoring models best represent the data. These improvements in creating ensembles of independent models will address the issue of how *precisely* cryo-EM data can be represented by an individual model. However, as in high-resolution crystallography (Smith 1986, Lang 2010), we anticipate there will be cases where simple model types will not suffice as accurate descriptions of discrete conformational heterogeneity. Time-averaged ensembles (Burnley 2012) and multiconformer models (Keedy 2015) represent two possible paths towards modeling the alternative conformations that underlie the spatially distinct, yet ensemble-averaged, density. While the analysis of patterns of clashes (van den Bedem 2013), time-resolved conformational changes (Cho 2010, Hekstra 2016), or temperature responses (Schmidt 2013, Keedy 2015) can be used to infer the coupling between different conformationally heterogeneous degrees of freedom, there is no direct route to deriving and validating this information from a single X-ray diffraction experiment. In contrast, the single particle nature of cryo-EM allows for focused reclassification to resolve the ensemble averaging of conformational heterogeneity into representative conformational states, and therefore the degree of long-range conformational coupling, across the entire macromolecule (Dashti 2014, des Georges 2016, Fernandez-Leiro and Scheres 2016). The limits of these approaches have not yet been identified, but it is reasonable to expect that they will progress to routine reclassification of rigid body motions of subdomains (Maji 2017) and perhaps secondary structures or even rotamers in the near future as instrumentation, data collection strategies, and processing algorithms improve. Evaluation of synthetic datasets, as used here to assess precision in modeling, will become increasingly important as the physical limits of reclassification are approached and the focus turns to validation and avoiding model bias.

The methodology outlined in this manuscript can be performed in an automated fashion, and the multiple models for every EMDB included in this study, as well as the per-residue analyses have been made available online (www.lander-lab.com/convergence). New depositions to the EMDB will be automatically downloaded and analyzed using the pipeline described here, and the results made available on the webserver. Furthermore, we have made the multi-resolution 20S and β-galactosidase density suites available for download at this site. Any future updates to the processing strategy for generating per-residue RMSDs will be announced on this website.

## Methods

### 20S proteasome processing

Super-resolution 8k × 8k micrograph movies (196 total movies, EMPIAR entry 10025 (Campbell 2015)) with a super-resolution pixel size of 0.655 Å were corrected for beam-induced motion using MotionCorr 2.0, a modified version of MotionCorr (Li 2013), using a 3 frame running average window. The resulting motion-corrected, summed frames were used for CTF estimation using CTFFind4 (Rohou and Grigorieff 2015). Any micrographs with a CTF confidence value below 0.95 were discarded. A difference of Gaussian (DoG) picker (Voss 2009) was used to select particles from the first 10 micrographs to yield an initial dataset of 627 particles. These particles were binned by 4 (2.62 Å/pixel and a box size of 128 pixels) and subjected to reference-free 2D classification using a topology representing network analysis (Ogura 2003) and multi-reference alignment in Appion (Lander 2009). The best 4 classes representing approximately all views were then used as templates against the entire dataset using FindEM (Roseman 2004) yielding 141,988 particles. 2x decimated particles (1.31 Å/pixel and a box size of 256 pixels) were then subjected to reference-free 2D-classification using RELION v1.4 (Scheres 2012). Particles (114,457) from the best classes were then subjected to 3D auto-refinement (0.655 Å/pixel and a box size of 512 pixels) using EMD-6287 (Campbell 2015) as an initial model (low-pass filtered to 60 Å) with D7 symmetry imposed. Subsequent movie-refinement (with a running average of 7 frames) was followed by particle polishing (using a standard deviation of 200 pixels on the inter-particle distance and a 3 frame B-factor running average (Scheres 2014)). The shiny particles were refined and subjected to a no alignment clustering using a mask against the full particle. This mask was generated using a 10-Å low-pass filtered version of the reconstructed map with a three-pixel extension and a five-pixel wide cosine-shaped soft edge. The best class (94,794 particles) from the resulting 3 classes was further refined using the same mask. The final resolution was estimated to ~2.7 Å using a gold-standard FSC cutoff of 0.143 (Rosenthal and Henderson 2003, Henderson 2012, Scheres and Chen 2012) after using phase-randomization to account for the convolution effects of a solvent mask on the FSC between the two independently refined half maps (Chen 2013) (**Figure 1–figure supplement 1**).

### β-galactosidase processing

Super-resolution 8k × 8k micrograph movies (EMPIAR entry 10061 (Bartesaghi 2015)) were downscaled 2x (yielding a final pixel size of 0.637 Å). Beam-induced motion correction and CTF estimation were performed as described for the 20S dataset. Any micrographs with a CTF confidence value below 0.95 were eliminated from further processing. DoG picker (Voss 2009) was used to select particles from the first 100 micrographs to yield an initial dataset of 12,195 particles. The best 8 classes representing approximately all views were then used as templates against the entire dataset using FindEM (Roseman 2004) yielding an initial data set of 140,393 particles. 4x downscaled particles (2.548 Å/pixel and a box size of 96 pixels) were then subjected to reference-free 2D-classification using RELION. 97,188 particles from the best classes were then subjected to 3D auto-refinement using EMD-2984 (Bartesaghi 2015) low-pass filtered to 60 Å as an initial model, with D2 symmetry imposed during the refinement. Particles were then re-centered and re-extracted, and all subsequent calculations were performed without binning (0.637 Å/pixel and a box size of 384 pixels). Movie-refinement with a running average of 7 frames was followed by particle polishing using a standard deviation of 1000 pixels on the inter-particle distance and a 3 frame B-factor running average (Scheres 2014). The shiny particles were refined and subjected to a no alignment clustering using a mask against the full particle. This mask was generated using a 15-Å low-pass filtered version of the reconstructed map with a three-pixel extension and a five-pixel wide cosine-shaped soft edge. The best class (71,379 particles) from the resulting 3 classes was further refined using the same mask. The final resolution was estimated to ~2.2 Å based on the gold-standard FSC 0.143 cutoff (Henderson 2012, Scheres and Chen 2012) (**Figure 1–figure supplement 9**).

### Lower resolution EM density generation

The following protocol was used to generate lower resolution structures (nominal FSC-reported value to ~5 Å in increments of ~0.2 Å) for both the 20S proteasome and β-galactosidase using particles from the refinements detailed above. For each structure, a separate RELION *.star* file was created (i.e. *20S_half1.star*) that contained the particles from each half map utilized during the gold-standard refinement that yielded the highest resolution structure. Each of the two star files was then back projected with each particle being subjected to a random translational offset (i.e. 0 - 3.1 sigma pixel error) using the *relion_reconstruct* command. For example, the following command was used to generate a half map of the 20S core particle with a translational offset of 1 sigma:

### relion_reconstruct ‐‐i 20S_half1.star ‐‐o 20S_1sigma_half1_class001_unfil.mrc ‐‐angpix 0.655 ‐‐sym D7 ‐‐ctf true ‐‐j 16 ‐‐shift_error 1

After generating the two half volumes for a desired translation error, the post-processing function within RELION was then used to create a single volume that had been sharpened and low-pass filtered (FSC-weighted) based on the FSC curve between the two half maps. The amount of per-particle translational shifts was adjusted empirically to obtain to a reconstruction at the approximate desired resolution. In total, 15 structures of the 20S proteasome from ~2.7 Å to ~4.9 Å resolution and 14 structures of β-galactosidase from ~2.2 Å to ~4.9 Å resolution were generated. Initially, both translational and rotational offsets were applied to generate lower resolution structures, but the resulting densities showed severe anisotropic loss of resolution. Adding even a small angular offset of 0.1 sigma resulted in reconstructions showing a severe loss of density in the peripheral regions, with only a small region of the core exhibiting structural features that corresponded to the FSC-reported resolution. For this reason, only translational offsets were applied in generating the multi-resolution suites.

### Multi-model pipeline

All Electron Microscopy Data Bank (EMDB, www.emdatabank.org) depositions generated using C- or D-symmetries (i.e. C1, C2, D7, etc., not icosahedral or helical symmetries) having a reported resolution better than 5 Å (FSC gold-standard) with an associated PDB were used in this study. If the deposited model for a symmetrized map only contained a single ASU, symmetry mates of the ASU were generated for these analyses. All EM densities were used without further modification. Initial models were generated by stripping PDB files of all alternate conformations, non-protein ligands/cofactors, resetting all occupancies to 1, and setting the isotropic B-factor to the approximate mean value. For entry 6287, model generation was performed as described in Campbell et al. 2015 prior to analysis using the pipeline detailed herein. Each entry was then refined using Rosetta with a selected output of 100 models using weighting terms based on reported resolution. The reported symmetry imposed during map generation was using during refinement. The 10 Rosetta-refined models that consistently scored best in categories such as geometry outliers (%, lower better), Ramachandran outliers (%, lower better), MolProbity clashscore (Chen 2010) (value, lower better), and Rosetta aggregate score (Wang 2016) (value, lower better) were selected as the “best” models for real-space refinement using the Phenix suite (Adams 2010) with NCS parameters determined by map symmetry. The reported resolution of the corresponding EM density was used for both refinement programs.

## Local resolution estimation and display

The ‘blocres’ function in the Bsoft package (Heymann and Belnap 2007) was used to generate local resolution maps, based on the two half-volumes outputted from the RELION 3D auto-refinement. All display images were generated using UCSF Chimera (Goddard 2007).

## Acknowledgements

We thank Jean-Christophe Ducom at The Scripps Research Institute High Performance Computing for computational support, and Yiru Xu for help performing preliminary analyses of low-resolution densities for the “resolution devolution”. We would also like to thank members of the Lander lab and Elizabeth Villa for critical discussion of the manuscript. M.A.H. is supported by a Helen Hay Whitney Foundation postdoctoral fellowship. J.S.F is supported as a Searle Scholar, a Pew Scholar, a Packard Fellow, by the National Science Foundation (STC-1231306) and the National Institutes of Health (GM063210). G.C.L. is supported as a Searle Scholar, a Pew Scholar, and by the National Institutes of Health (DP2EB020402).

